## Supplementary materials for "Delayed induction of type I and III interferons mediates nasal epithelial cell permissiveness to SARS-CoV-2"

### Contents

Supplementary Figures S1-10

Supplementary Tables S1-6

Supplementary Methods

Note supplementary datasets S1-5 (csv files) are not included in this file

**Supplementary Figures**

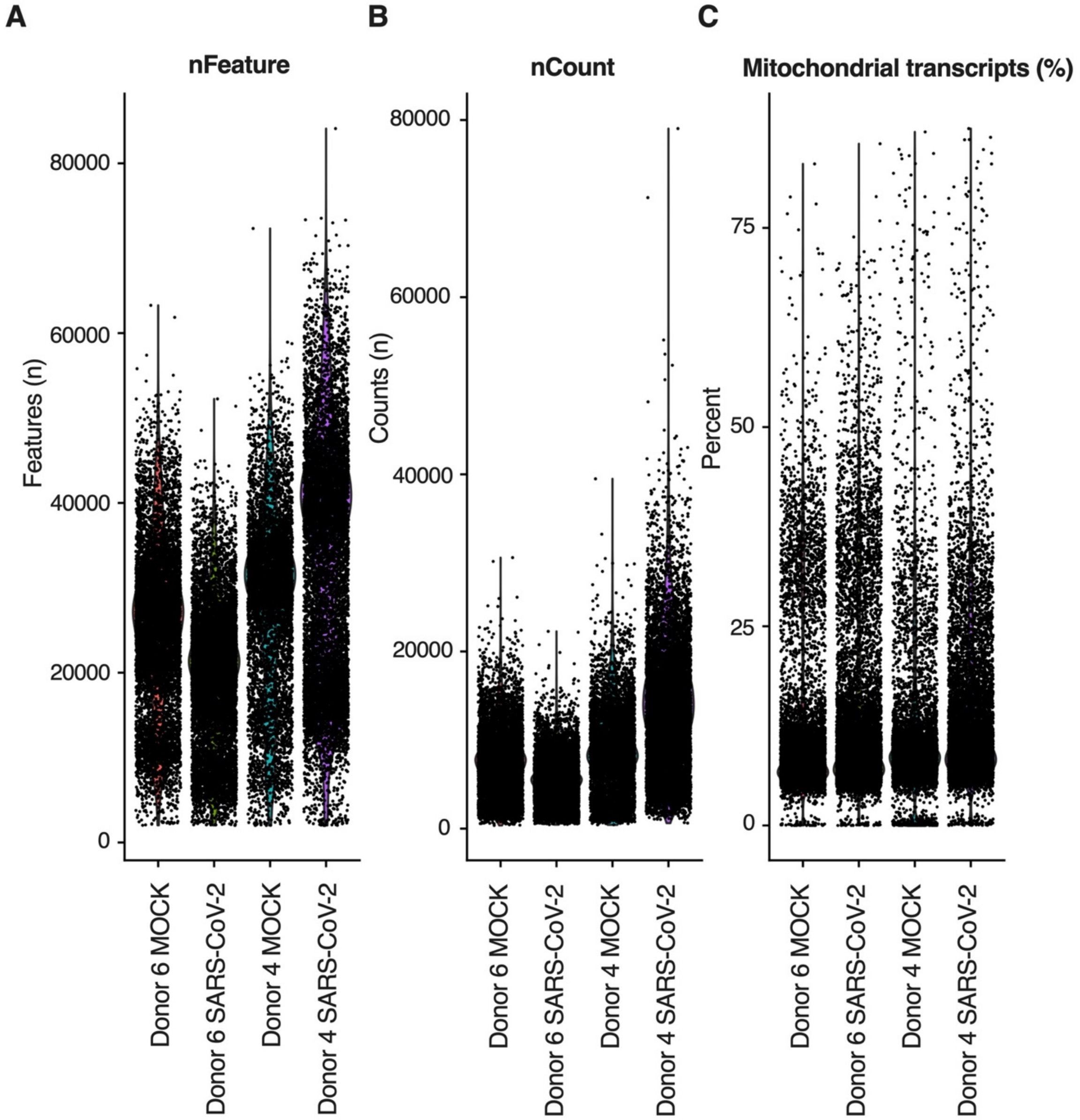


**Figure S1.** Single cell RNA sequencing quality control plots. Violin plots, split by sample, showing (A) the total number of genes detected in each cell (B) the total number of counts detected in each cell and (C) the proportion (as a percentage) of mitochondrial transcripts in each cell. For individual QC metrics see also Table S1.

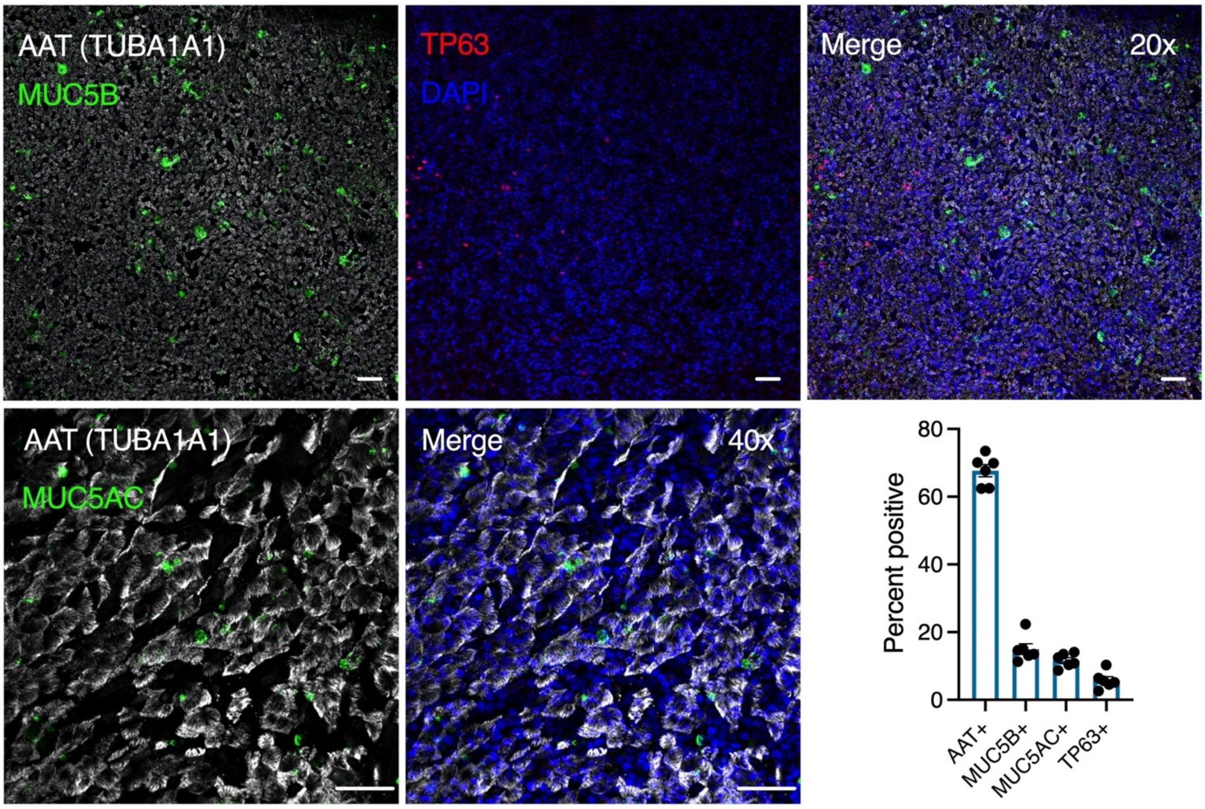


**Figure S2.** Immunofluorescence analysis of ciliated (AAT+), secretory (MUC5B+), goblet (MUC5AC+) and basal (TP63+) cells in nasal ALI cultures. Representative images from n=6 donors, scale bar = 20 mm (top panel 20x magnification, bottom panel 40x magnification as indicated). Frequency of cell type as a proportion of cells counted is displayed in the bar plot. MUC5B and MUC5AC co-staining demonstrated no overlap in immunoreactivity (data not shown).

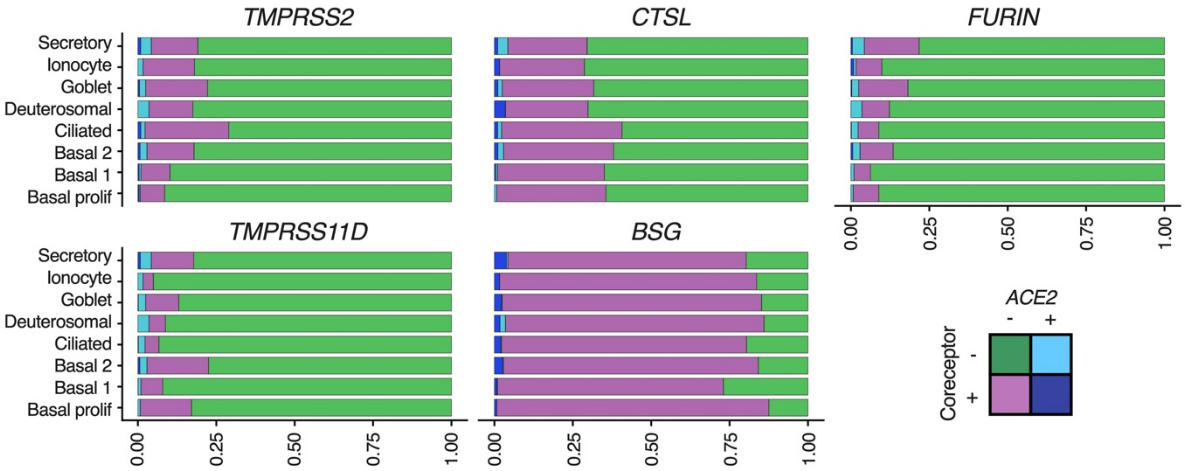


**Figure S3.** Single-cell RNA-seq analysis of entry receptor expression by cell type. Bars represent the proportion of cells expressing each combination of *ACE2* and other transcript, coloured according to the key. Dark blue represents the proportion of cells of each type expressing both *ACE2* and the relevant additional transcript. Data from analysis of 28,346 cells from n=2 donors.

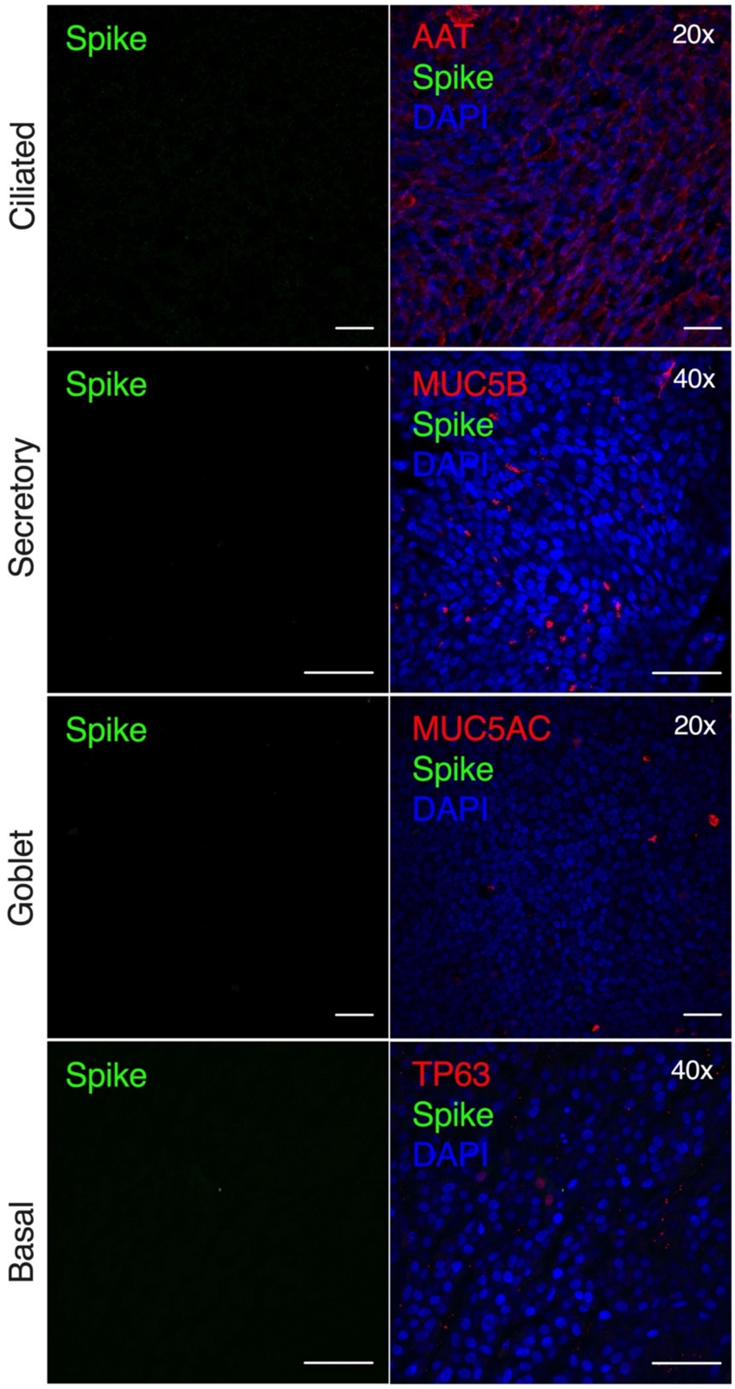


**Figure S4.** Immunofluorescence analysis of S protein immunoreactivity in mock infected nasal ALI cultures. Displayed are mock infected ciliated (AAT+), secretory (MUC5B+), goblet (MUC5AC+) and basal (TP63+) cells stained for S protein. Representative images from n=5 donors, scale bar = 20 mm (images at 20x or 40x magnification as indicated).

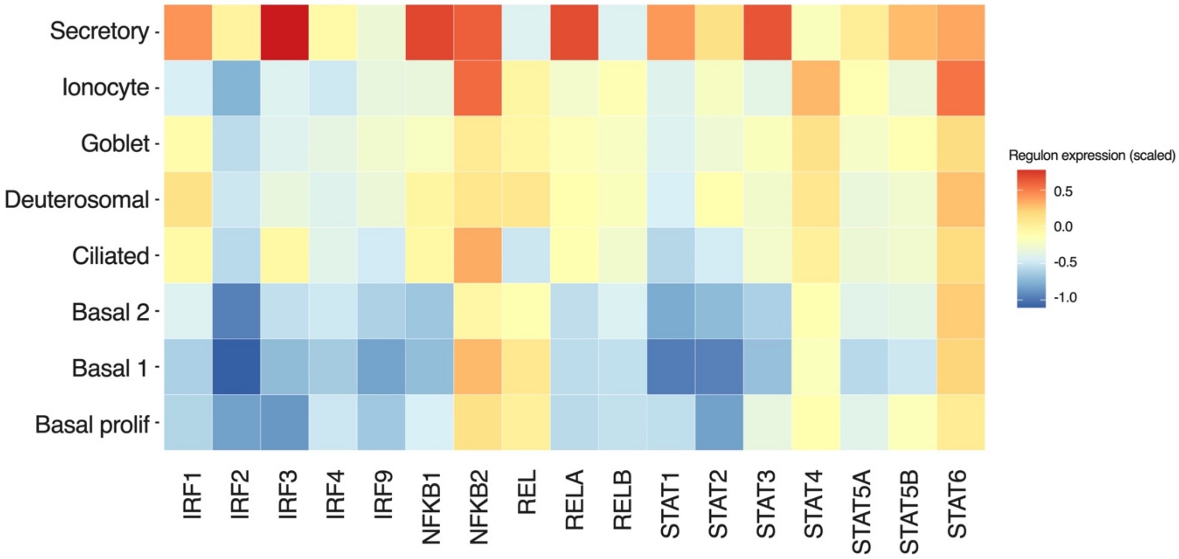


**Figure S5.** DoRoTHea/VIPER analysis of regulon activity in infected cells. Median regulon activity per cluster in infected cells, corrected for activity in uninfected cells by subtraction then Z-normalised by TF (i.e. values > 0 imply TF more active in infected cells). Data from analysis of 28,346 cells total to estimate regulon activity of which 8,861 infected, from n=2 donors at 24 hpi.

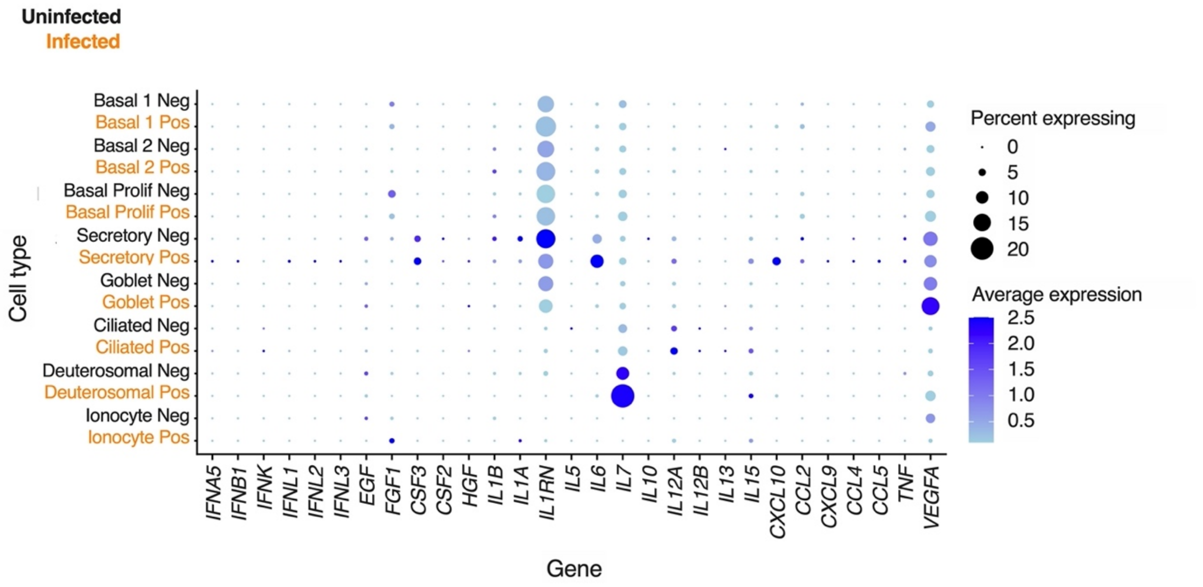


**Figure S6.** Interferon, chemokine and cytokine induction in response to SARS-CoV-2. Dot plot showing single-cell RNA-seq analysis of cytokine and chemokine transcript detection in n=2 donors at 24 hpi (size of dots represent proportion of cells expressing and colour represents mean expression). Uninfected cells are labelled black (Neg) and infected cells orange (Pos) based on expression of SARS-CoV-2 mRNA. Low-level induction of certain proinflammatory cytokines (*IL6, IL12A, IL15*), chemokines (*CXCL9, CXCL10*) and *VEGFA* is demonstrated in SARS-CoV-2-infected nasal cells.

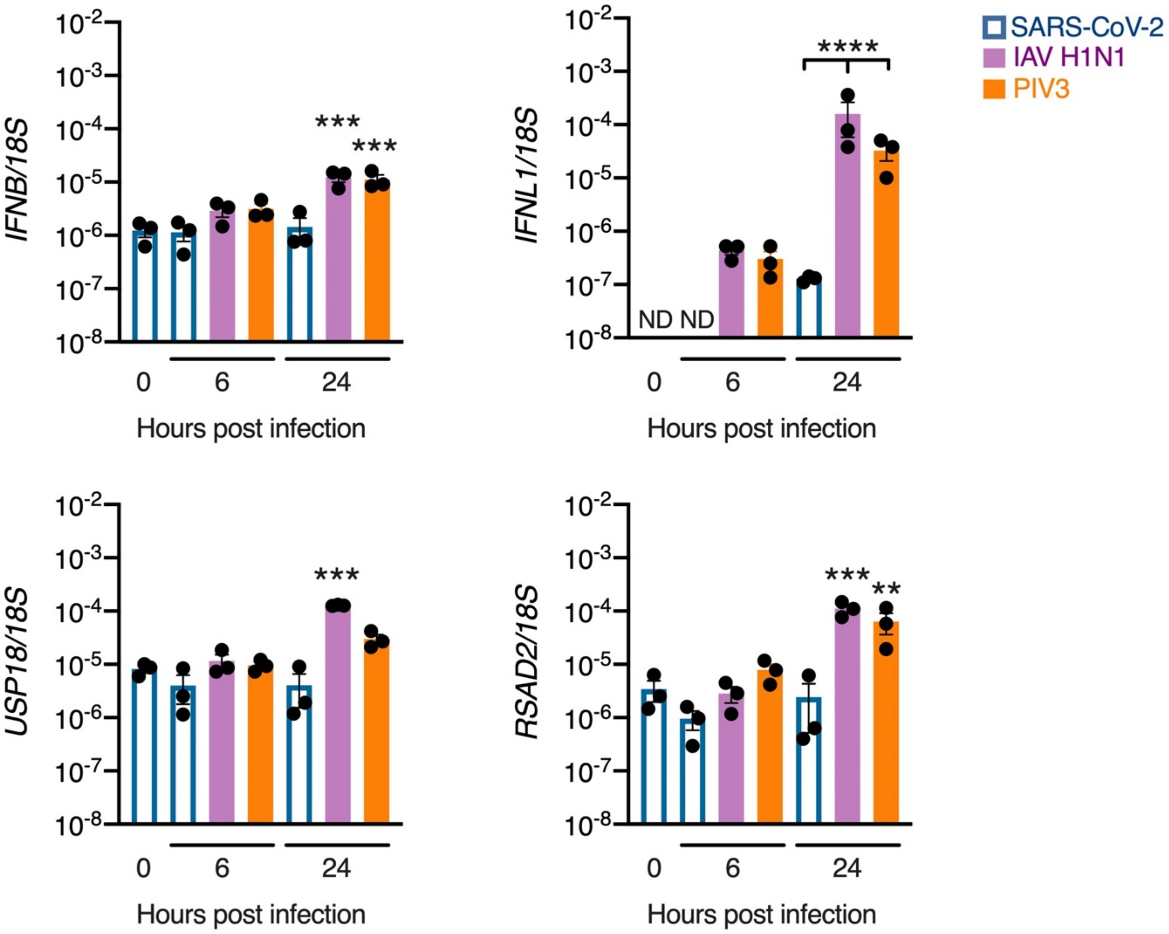


**Figure S7**. Delayed induction of IFNs and ISGs in response to SARS-CoV-2 compared to other viruses. RT-PCR analysis of *IFNB*, *IFNL1*, *USP18* and *RSAD2* expression in nasal ALI cultures mock infected (0h) or exposed to SARS-CoV-2 (open bars), influenza A virus (IAV H1N1, purple bars) or parainfluenza 3 virus (PIV3, orange bars) for the times displayed, all at MOI 0.1 (n=3 donors, mean ± SEM; ANOVA with Dunnett's post-test correction compared to 0h, or Sidak's post-test correction [all viruses compared at 24 hpi], ** P < 0.01 *** P < 0.01 **** P < 0.001). ND = not detected.

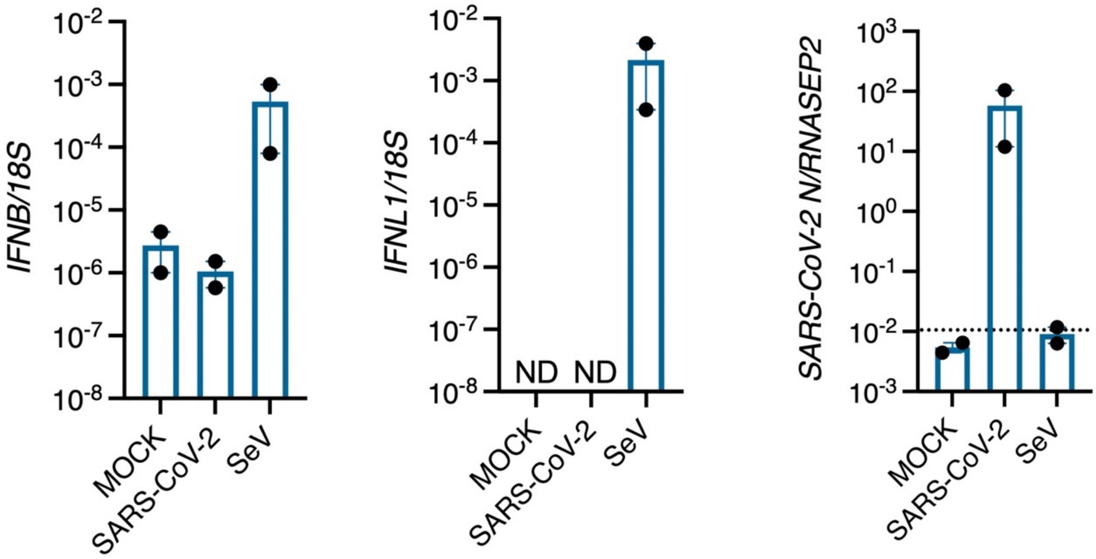


**Figure S8**. Robust nasal cell expression of *IFNB* and *IFNL1* in response to Sendai virus. RT-PCR analysis of *IFNB*, *IFNL1* and SARS-CoV-2 *N* gene expression in nasal ALI cultures exposed to SARS-CoV-2 (MOI 2) or a DVG-rich stock of Sendai virus (SeV) for 6 h (n=2 donors, mean ± SEM). ND = not detected.

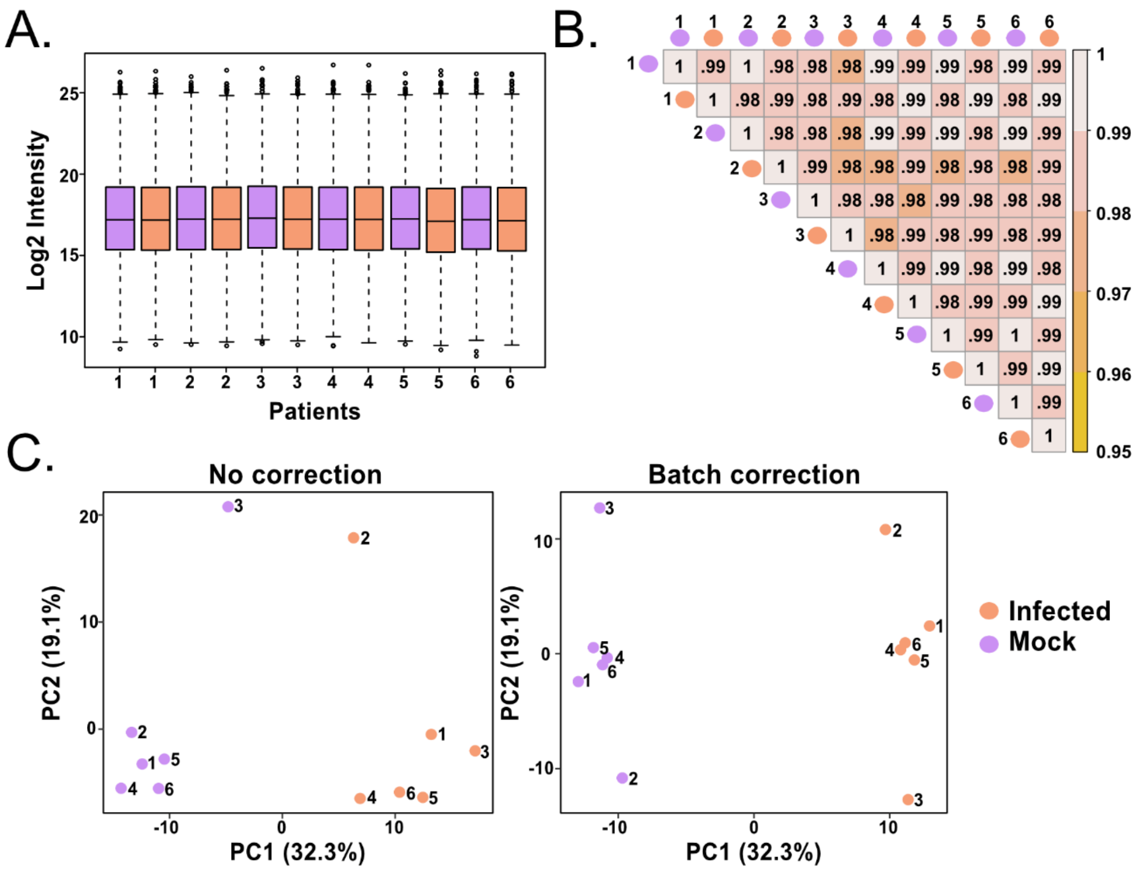


**Figure S9.** Quality control measures for the proteomics data set. (A) Boxplot of log2 transformed samples shows equal loading. (B) Pearson correlation heatmap among the log2 transformed samples shows high reproducibility between samples. (C) Principal component analysis plots with no correction (left) and after removing patient batch effects (right).

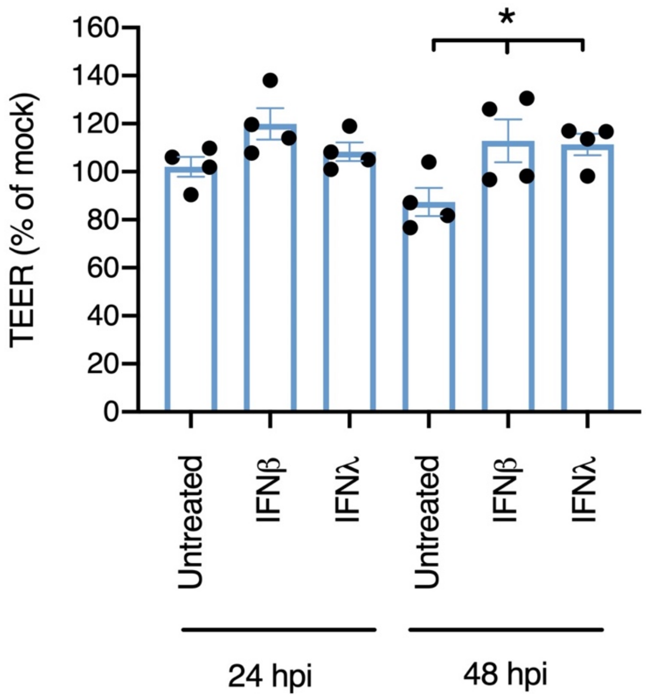


**Figure S10.** IFN treatment preserves barrier integrity in the face of SARS-CoV-2 infection. Trans-epithelial resistance measurement (expressed as % of mock infected controls) at 24 and 48 hpi (MOI 0.01) were compared to cells pre-treated for 16h with IFNβ1 (1000 IU/mL) or IFNλ1 (100 ng/mL). Repeat experiments in n=4 donors, mean ± SEM; * $P < 0.05$, ANOVA with Sidak's post-test correction.

### Supplementary Tables

| Sample_id | Total number of reads | Mean reads per cell | Alignment rate (%) | Reads mapped to GRCh38 (5) | Reads mapped to SARS-CoV-2 | Estimated number of cells |
|---|---|---|---|---|---|---|
| Donor4_Mock | 184,897,338 | 14,329 | 91.3 | 91.3 | 0 | 12,904 |
| Donor4_Infected | 807,865,486 | 61,100 | 87.7 | 77.1 | 10.8 | 13,222 |
| Donor6_Mock | 232,499,890 | 17,121 | 90.4 | 90.4 | 0 | 13,580 |
| Donor6_Infected | 174,974,839 | 13,653 | 90 | 88.8 | 1.3 | 12,816 |

**Table S1.** Single cell RNA-seq post-alignment quality control metrics. Quality control output from CellRanger following alignment.

| GO_TERM | FDR Adjusted P value |
|---|---|
| Defence response to virus | 4.5E-28 |
| Type I interferon signalling pathway | 2.3E-18 |
| Response to virus | 1.4E-13 |
| Negative regulation of viral genome regulation | 5.7E-11 |
| Interferon gamma-mediated signalling pathway | 5.8E-9 |
| Innate immune response | 7.6E-5 |
| Intracellular transport of viral protein in host cell | 6.9E-3 |
| Negative regulation of type I interferon production | 7.2E-3 |
| Antigen processing and presentation via MHC class I | 1.1E-2 |
| Cellular response to interferon alpha | 1.7E-2 |
| Response to interferon alpha | 2.0E-2 |

**Table S2.** Pathway analysis of proteomics data showing upregulated pathways. Displayed are pathways with Benjamini-Hochberg false-discovery rate (FDR)-adjusted P value < 0.05 (5E-2).

| GO_TERM | FDR Adjusted P value |
|---|---|
| TRIF-dependent toll-like receptor signalling pathway | 1.5E-2 |
| Regulation of transcription from RNA polymerase II promoter in response to hypoxia | 1.5E-2 |
| Endosomal transport | 1.7E-2 |
| MyD88-independent toll-like receptor signalling pathway | 1.7E-2 |
| Transcription-coupled nucleotide-excision repair | 2.4E-2 |

**Table S3.** Gene Ontology (GO) Term analysis of proteomics data showing downregulated pathways. Displayed are pathways with Benjamini-Hochberg false-discovery rate (FDR)-adjusted P value < 0.05 (5E-2).

| Donor no. | Sex | Age (years) |
|---|---|---|
| 1 | Female | 46 |
| 2 | Female | 38 |
| 3 | Male | 68 |
| 4 | Male | 78 |
| 5 | Female | 54 |
| 6 | Male | 41 |

**Table S4.** Nasal cell donors.

| Gene | UPL probe | Forward sequence | Reverse sequence |
|---|---|---|---|
| *IFNB* | #25 | CGACACTGTTCGTGTTGTCA | GAAGCACAACAGGAGAGCAA |
| *IFNL1* | #75 | GGGACCTGAGGCTTCTCC | CCAGGACCTTCAGCGTCA |
| *IL6* | #40 | GATGAGTACAAAAGTCCTGATCCA | CTGCAGCCACTGGTTCTGT |
| *IL1B* | #78 | TACCTGTCCTGCGTGTTGAA | TCTTTGGGTAATTTTTGGGATCT |
| *RSAD2* | #9 | GAGGGTGAGAATTGTGGAGAAG | GCGCTCCAAGAATCTTTCAA |
| *USP18* | #44 | CAACGTGCCCTTGTTTGTC | ATCAGGTTCCAGAGTTTGAGGT |
| *ISG15* | #23 | GCGAACTCATCTTTGCCAGTA | CCAGCATCTTCACCGTCAG |
| *18S* | #81 | CCGATTGGATGGTTTAGTGAG | AGTTCGACCGTCTTCTCAGC |

**Table S5.** Primers/probes. UPL = Roche universal probe library.

| Antibody | Host | Dilution | Source | Code |
|---|---|---|---|---|
| Spike | Rabbit | 1:1000 | Novus | nb100-56578 |
| RSAD2 | Rabbit | 1:1000 | CST | 13996 |
| ISG15 | Rabbit | 1:1000 | CST | 2743 |
| USP18 | Mouse | 1:2000 | SCB | sc-1668 |
| ACE2 | Rabbit | 1:1000 | Abcam | ab15348 |
| ACE2 | Goat | 1:200 | R&D | AF933 |
| TMPRSS2 | Rabbit | 1:1000 | Abcam | ab92323 |
| MxA | Rabbit | 1:1000 | SCB | sc-50509 |
| GAPDH | Rabbit | 1:10,000 | CST | 5174 |
| MUC5B | Rabbit | 1:1000 | Sigma | HPA008246 |
| MUC5AC | Rabbit | 1:1000 | Sigma | HPA040615 |
| TP63 | Mouse | 1:2000 | Abcam | ab735 |
| Acetylated-alpha tubulin | Mouse | 1:1000 | Abcam | ab24610 |
| Anti-rabbit HRP- conjugated | Goat | Primary-dependent | CST | 7074 |
| Anti-mouse HRP-conjugated | Horse | Primary-dependent | CST | 7076 |
| AF488 conjugated anti-mouse | Goat | 1:2000 | TFS | A-11001 |
| AF488 conjugated anti-rabbit | Goat | 1:2000 | TFS | A-11008 |
| AF594 conjugated anti-mouse | Goat | 1:2000 | TFS | A-11005 |
| AF594 conjugated anti-rabbit | Goat | 1:2000 | TFS | A-11012 |

**Table S6**. Antibodies. CST = Cell Signalling; SCB = Santa Cruz Biotechnology; R&D = R&D biosystems; TFS = ThermoFisher Scientific. HRP = horseradish peroxidase.

### Supplementary Methods

#### Proteome sample preparation

Cells were washed three times with cold PBS before addition of solubilisation buffer (5% (w/v) SDS, 50 mM TEAB) to the apical compartment for 10 min at room temperature (RT). Samples were heated at 75°C for 45 min, before freezing and stored at -80°C. Protein concentration was determined by EZQ® protein quantification assay. A total of 30 µg protein was reduced by incubation with 5 mM tris(2-carboxyethyl)phosphine for 15 min at 37°C, and subsequently alkylated with 20 mM iodoacetamide for 30 min at RT in the dark. Protein digestion was performed using the suspension trapping (S-Trap™) sample preparation method according to the manufacturer's guidelines (ProtiFi, USA). Briefly, 2.5 µL of 12% phosphoric acid was added to each sample, followed by the addition of 165 µl S-Trap binding buffer (100 mM TEAB in 90% methanol, pH 7.1). Samples were added to S-Trap Micro spin columns followed by centrifugation (4,000 g, 2 min). Each S-Trap Mini-spin column was washed with 150 µL S-trap binding buffer by centrifugation (4,000 g, 1 min). This process was repeated for a total of 4 washes. 25 µL of 50 mM TEAB, pH 8.0 containing trypsin (1:20 ratio of trypsin to protein) was added to each sample, followed by proteolytic digestion for 3 hours at 47°C without shaking. Peptides were eluted with 50 mM TEAB pH 8.0 and centrifugation (4,000 g, 2 min). Elution steps were repeated twice more, using 0.2% formic acid and 0.2% formic acid in 50% acetonitrile, respectively. The three eluates from each sample were combined and dried using a speed-vac before storage at -80°C.

#### TMT-16 plex labelling

Each 30 µg protein digest was resuspended in 25 µL 100 mM HEPES, pH 8.5. TMT-16 plex labelling (TMT lot number: UI292951) was carried out as per the manufacturer's instructions. Samples were assigned to a TMT tag. 10 µL of the corresponding TMT tag was added per sample and incubated for 1 hour at RT. An aliquot corresponding to 1 µg was taken from each sample and pooled together for ratio and labelling efficiency checks, prior to making the full pooled sample. The test pool was quenched with 0.69 µL of 5% hydroxylamine, incubated for 15 min at room temperature, and dried using a speed-vac. The sample was cleaned using a C18 spin column as per the manufacturer's guidelines (Thermo Scientific), and subsequently dried using a speed-vac. Peptides (dissolved in 5% formic acid) from the pooled sample were analysed for labelling efficiency and ratio check. For the ratio check, each sample (corresponding to a single TMT channel) was normalised to the average summed intensity of all samples within its pool. Each sample was quenched with 2.5 µL 5% hydroxylamine and incubated for 15 min. Subsequently, samples were pooled together based on the scaling factors, which were calculated using the test pool. Samples were dried using a speed-vac, cleaned using MacroSpin columns as per the manufacturer's guidelines (Harvard Apparatus, USA), and dried down again using a speed-vac prior to offline high-performance liquid chromatography (HPLC) fractionation.

**Offline HPLC Fractionation**

Peptides were resuspended in 80 µL ammonium formate, pH 8.0. Peptides were fractionated on a Basic Reverse Phase column (Gemini C18, 3 um particle size, 110A pore, 3 mm internal diameter, 250 mm length, Phenomenex #00G-4439-Y0) on a Dionex Ultimate 3000 off-line LC system. All solvents used were HPLC grade (Rathburn Chemicals, UK). 40 µL of peptide sample were loaded onto the column for 1 min at 250 µL/min using 99% Buffer A (20 mM ammonium formate, pH 8.0) and eluted for 40 min on a linear gradient from 1 to 90% Buffer B (100% acetonitrile (ACN)). Peptide elution was monitored by UV detection at 214 nm. Fractions were collected every minute from 2 to 38 minutes for a total of 36 fractions. Fractions were pooled using non-consecutive concatenation to obtain 18 pooled fractions (e.g. pooled fraction 1: fraction 1 + 19). Each fraction was acidified to a final concentration of 1% TFA and dried using a speed-vac.

**Mass spectrometry**

Peptides were dissolved in 5% formic acid, and each sample was independently analysed on an Orbitrap Fusion Lumos Tribrid mass spectrometer (Thermo Fisher Scientific), connected to an UltiMate 3000 RSLCnano System (Thermo Fisher Scientific). Peptides (~2 µg per fraction) were injected on an Acclaim PepMap 100 C18 LC trap column (100 µm ID × 20 mm, 3 µm, 100 Å) followed by separation on an EASY-Spray nanoLC C18 column (75 ID µm × 750 mm, 2 µm, 100 Å) at a flow rate of 200 nL/min. Solvent A was 0.1% formic acid in $H_2O$ and solvent B was 80% ACN containing 0.1% formic acid. The gradient used for analysis of proteome samples was as follows: solvent B was maintained at 3% for 5 min, followed by an increase of solvent B from 3% to 35% in 120 min, 35% to 90% B in 0.5 min, maintained at 90% B for 4 min, followed by a decrease to 3% in 0.5 min and equilibration at 3% for 20 min. Mass spectrometric identification and quantification was performed on an Orbitrap Fusion Tribrid mass spectrometer (Thermo-Fisher Scientific) operated in data-dependent, positive ion mode. Full scan spectra were acquired in a range from 375 m/z to 1500 m/z, at a resolution of 120,000, with a standard automated gain control (AGC) (Tune 3.3) and a maximum injection time of 50 ms. Precursor ions were isolated with a quadrupole mass filter width of 0.7 m/z and CID fragmentation was performed in one-step collision energy of 30% and 0.25 activation Q. Detection of MS/MS fragments was acquired in the linear ion trap in a rapid mode using a Top 3s method, with a standard AGC target and a maximum injection time of 50 ms. The dynamic exclusion of previously acquired precursor was enabled for 60 s with a tolerance of +/-10 ppm. Quantitative analysis of TMT-tagged peptides was performed using FTMS3 acquisition in the Orbitrap mass analyser operated at 60,000 resolution, with a standard AGC target and maximum injection time of 118 ms. HCD fragmentation on MS/MS fragments was performed in one-step collision energy of 55% to ensure maximal TMT reporter ion yield and synchronous-precursor-selection (SPS) was enabled to include 10 MS/MS fragment ions in the FTMS3 scan.

**Mass spectrometry data analysis**

All spectra were analysed using MaxQuant 1.6.10.43 and searched against SwissProt *Homo sapiens* (with 42423 sequences) and Trembl SARS-CoV-2 (with 107 sequences) FASTA files. Peak list generation was performed within MaxQuant and searches were performed using default parameters and the built-in Andromeda search engine. Reporter ion MS3 was used for quantification and the additional parameter of quantitation labels with 16 plex TMT on N-terminus or lysine was included. The enzyme specificity was set to consider fully tryptic peptides, and two missed cleavages were allowed. Oxidation of methionine and N-terminal acetylation were allowed as variable modifications. Carbamidomethylation of cysteine was allowed as a fixed modification. A protein and peptide false discovery rate (FDR) of less than 1% was employed in MaxQuant. Reporter ion intensities were used for data analysis. Briefly, the data were filtered to remove proteins that matched to a contaminant or a reverse database, which were only identified by site, which were not quantified in every sample, or which contained less than 2 unique peptides. Reporter ion intensity values were $\log_2$ transformed. Each sample within a TMT set was then normalised to the average median intensity of all 12 samples within that set. Moderated *t*-tests, with patient accounted for in the linear model, was performed using Limma, where proteins with an adjusted $P < 0.05$ were considered as statistically significant. Proteins with differential abundance (adjusted p-value <0.05 and fold change > 1.5) were analysed using the search tool for retrieval of interacting genes (STRING) database version 11 (https://string-db.org/). The data was modified for presentation using Cytoscape version 3.7.2. Proteins were grouped by functional categories based Uniprot annotation (https://www.uniprot.org). Active interaction sources, including experiments and databases, and an interaction score > 0.7 were applied to construct the protein-protein interaction networks. In the network, the nodes correspond to the proteins identified and the edges represent the interactions. The node colour gradient depicts fold change in protein expression in infected compared to mock samples. All analysis was performed using R 3.6.2.
